# Tracing mobility among Eneolithic-Bronze Age Kurgan populations in the North Pontic steppe

**DOI:** 10.64898/2026.03.21.713323

**Authors:** Alexey G. Nikitin, Virginie Renson, Svitlana Ivanova, Nadia C. Neff, Haruan Straioto, Sofiia Svyryd

**Affiliations:** Department of Biology, Grand Valley State University, Allendale, Michigan, United States of America; Archaeometry Laboratory, University of Missouri Research Reactor, Columbia, MO, United States of America; Institute of Archaeology, Ukrainian Academy of Sciences, Odesa, Ukraine; Department of Anthropology, University of New Mexico, Albuquerque, NM, United States of America; Museum of Archaeology and Ethnology, University of São Paulo, Brazil

## Abstract

Five millennia ago, nomadic people from the North Pontic steppe left a profound impact on the course of Eurasian prehistory. However, little is known about their mobility patterns within their home region. To address this knowledge gap, we conducted a survey of the strontium isotope landscape of people interred in the 4th-3rd millennium BCE burial mounds (kurgans) of the western part of the North Pontic steppe. By analyzing the strontium signature in human bone and dentin, we established strontium baseline values for the region. We subsequently correlated enamel strontium ratios from 25 selected individuals with the baseline obtained and with published strontium data across the North Pontic steppe. Enamel strontium ratios show that some individuals interred in the northwest North Pontic fall within the regional baseline range, whereas others overlap with values reported for the eastern North Pontic steppe. In conjunction with carbon (δ^13^C) and nitrogen (δ^15^N) stable isotope data, we further determined that some individuals interred in the western Pontic steppe either spent the later part of life in the west Caspian steppe or were affected by physiological stress during lifetime. By integrating our data with published isotopic datasets, we produced a first baseline heatmap of the North Pontic steppe for the c. 4000-2000 BCE chronological period.

## Introduction

In the Early Bronze Age (EBA, c. 3300-2200 BCE according to southeast European chronology), a wave of innovations swept across the Eurasian continent, introducing new linguistic features in the form of branches of Indo-European language, a developed pastoralist economy, and a hierarchal social and ritual system. These changes were accompanied by a change in genetic ancestry across Europe. The Anatolian Neolithic Farmer-based ancestry of early farming communities of Europe was replaced by a genetic package founded on the ancestry of Caucasus-Lower Volga migrants admixed with Neolithic North Pontic hunter-fisher-foragers [1]. The nomadic people of the North Pontic region carrying this genetic “steppe” package are referred to as the Kurgan people because they erected earth burial mounds throughout the steppe and adjacent forest-steppe [2]. Although their genetic profiles are well characterized, far less is understood about other aspects of their life history, including patterns of mobility within their home range. We set out to address this knowledge deficiency by conducting exploratory analysis of strontium isotope ratios from Eneolithic-EBA interments in kurgans in the west portion of the Pontic steppe (the Odesa region of Ukraine) focusing on a set of samples of high interpretive potential, to establish a foundation for future large-scale isotopic studies of mobility in the region from which the Early Bronze Age “steppization” of Europe was launched.

Strontium isotope analysis serves as a geochemical proxy for determining the geographic origins of individuals buried in the ground, providing valuable insights into patterns of mobility. When contextualized, this geochemical evidence can shed light on the social significance of burial inclusion or exclusion. For instance, comparing ^87^Sr/^86^Sr values from dental enamel with locally established baseline ranges can help distinguish between individuals from local and non-local backgrounds. The presence of both local and non-local signatures among the buried population suggests a community that embraced individuals from diverse geographic origins. In this context, “local” denotes conformity to the average bioavailable regional geographic zone, or, in our case, the isotopic compatibility with the regional geochemical landscape within which the mobile steppe groups likely circulate. Conversely, multiple non-local values may indicate a structured pattern of mobility.

The geological composition of the Odesa Region in Ukraine is a complex blend of Precambrian crystalline basement rocks and younger sedimentary layers (Fig 1). The region, part of the Black Sea Lowland, is influenced by the Black Sea Depression and the Scythian Plate. The Odesa region is located near the Podolian (Buh-Dniester) domain, which, along with the Azov domain, are the oldest components of the Ukrainian shield. The Cretaceous and Quaternary sedimentary fluvio-colluvial deposits in the area mostly consist of loess and loess-like loams. Pontian limestones from the Late Miocene (Upper Cenozoic) era are exposed in the coastal areas of the region [3–6].

**Fig 1.**
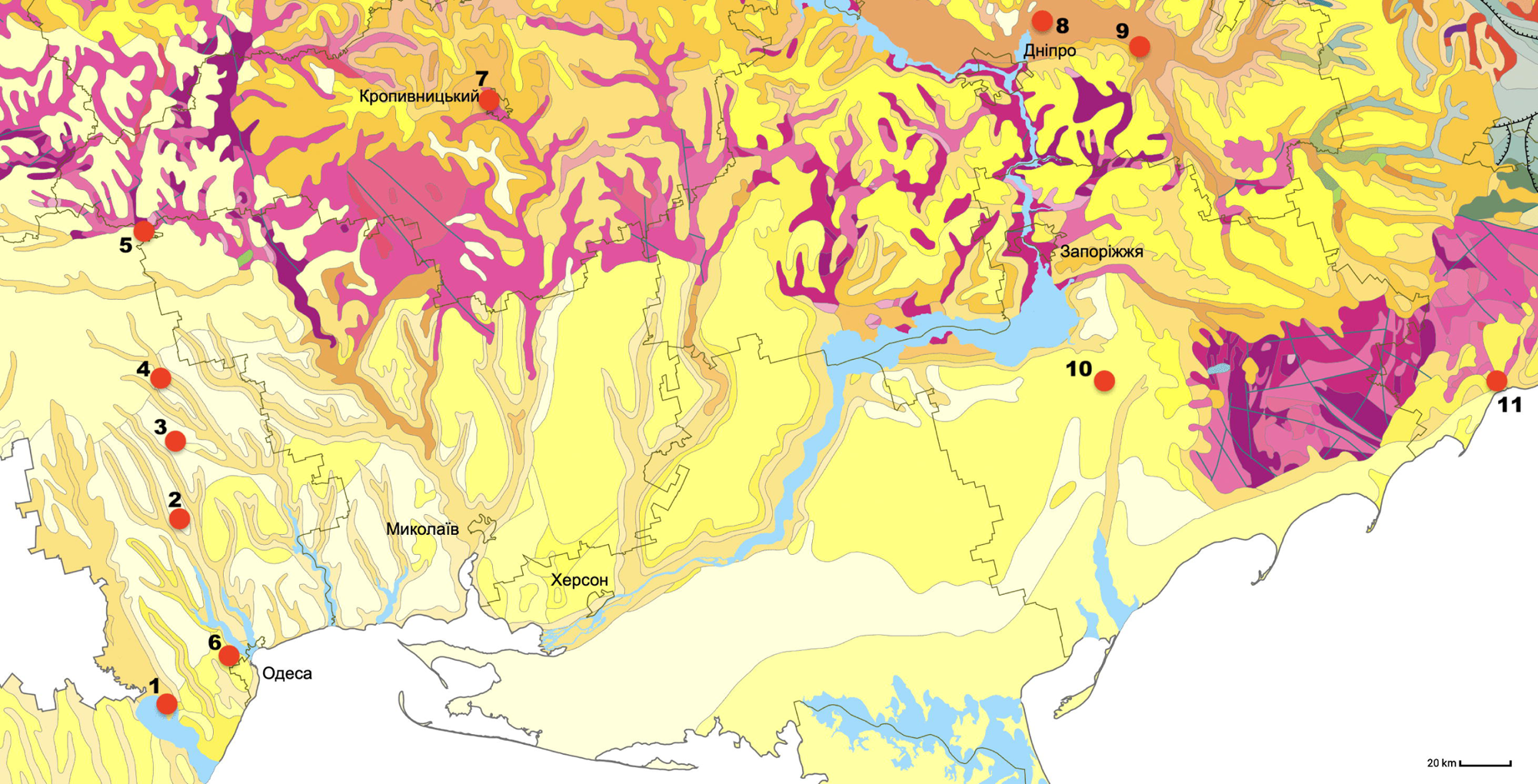
Geological map of the north Pontic steppe region showing kurgans from which samples were obtained for the present study and the sites used in the comparative analysis. 1, Mayaky; 2, Katarzhyno kurgans 1 and 2; 3, Revova kurgan 3; 4, Liubasha kurgan 2; 5, Dubynove kurgan 1; 6, Usatove-Velykyj Kuyalnik; 7, Sugokleya kurgan; 8, Pishchanka kurgan; 9, Shakhta Stepova kurgan group; 10, Vynohradne kurgan group; 11, Mariupol Neolithic necropolis. Sand, yellow and beige – Neogene, Pliocene; light and darker brown – Neogene, Miocene; darkest brown – Paleogene, Neocene; light pink – Archaean, Neoarchaean; dark pink – Archaean, Eoarchean; crimson – Proterozoic, Paleoproterozoic; green – Lower Cretaceous. Geological map by A. Grachev, geomap.land.kiev.ua, reproduced with permission. Basemap - Template_europe_map. Wikimedia Commons CC BY-SA.3.0.

Carbon (δ^13^C) and nitrogen (δ^15^N) isotope analysis of bone collagen offers a complementary perspective on an individual’s life history because these systems reflect the isotopic composition of mainly dietary protein, and secondarily, in varying proportions depending on nutritional status, carbohydrates, and lipids consumed over the last 7-20 years of life [7]. While ^87^Sr/^86^Sr ratios primarily indicate movement across different geological landscapes, δ^13^C and δ^15^N values can help differentiate individuals whose diets may have been shaped by different ecological zones, subsistence practices, or physiological conditions [8,9]. In highly mobile pastoral groups such as those of the North Pontic steppe, δ^13^C and δ^15^N values of collagen are therefore useful for assessing whether individuals with unusual strontium signatures also had different dietary histories later in life, which can strengthen interpretations of mobility and regional interaction.

In this study, we employ a multiproxy isotopic approach combining ^87^Sr/^86^Sr and δ^13^C and δ^15^N isotope values of bone collagen to investigate the life histories of individuals interred in seven Eneolithic-Bronze Age kurgans in the northwestern North Pontic region. To strengthen the reliability of mineral-based isotopic interpretations, we also evaluated the preservation of selected bioapatite samples using Attenuated Total Reflectance Fourier Transform Infrared Spectroscopy (ATR-FTIR) prior to ^87^Sr/^86^Sr analysis. FTIR is performed to screen bones or teeth for diagenetic alteration and confirm that the mineral structure is sufficiently preserved so the measured strontium ratios reflect the individual’s original biogenic signal rather than post-burial contamination. The analyzed specimens were selected to explore factors linking individuals within and between kurgans and to evaluate the extent to which such data can inform reconstructions of mobility across the North Pontic steppe during the 4^th^-3^rd^ millennium BCE, thereby guiding future large-scale sampling strategies. This research specifically investigates whether individuals buried in northwest North Pontic kurgans exhibit strontium isotope ratios consistent with locally derived baseline values or instead reflect mobility within or beyond the broader steppe throughout their lives. To address this, the study first establishes ^87^Sr/^86^Sr baseline values from the available dataset and then evaluates individual mobility patterns against this framework. This study further evaluates whether individual δ¹³C and δ¹⁵N values provide independent lines of evidence for mobility beyond that inferred from strontium isotope analysis alone. Ultimately, we integrate the newly obtained ⁸⁷Sr/⁸⁶Sr data with previously published results to establish the first regional isotopic baseline framework for the North Pontic region.

## Methods

### Provenance of archaeological samples

Samples detailed in this report came from a chain of kurgans in the Odesa Region, southern Ukraine (Fig 1). The Mayaky kurgan cemetery is situated on a promontory of the Dniester Estuary on the Black Sea coast, while the other five kurgans are located inland, to the north from Mayaky. Katarzhyno kurgans 1 and 2 were excavated in 1990-1991 by expeditions of the Odesa Archaeological Protection Center, which existed under the Ukrainian Society for the

Protection of Historical and Cultural Monuments. Dubynove 1, Liubasha 2 and Revova 3 kurgans were excavated in 2003 by an expedition of the Department of Archaeology of the North-Western Black Sea Region of the Institute of Archaeology of the National Academy of Sciences of Ukraine [10]. The kurgan cemetery at Mayaky was excavated by the Odesa Archaeological Museum of the National Academy of Sciences of Ukraine in 1974-1975, 1986, and 2009 [11–13]. The excavations were carried out under state permits and in accordance with Ukrainian heritage law. Following excavation, the remains were accessioned into the Odesa Archaeological Museum collection, where they are curated as part of the national archaeological archive. Detailed description of the sites and burials can be found in [1,10–14], and additional details are provided in S1 File.

Enamel, dentin, and cortical bone samples were obtained from 25 individuals representing Eneolithic Usatove (3800-3100 BCE), Steppe Eneolithic (3950-3300 BCE), EBA Yamna (3300-2500 BCE) and Catacomb (2800-2200 BCE), as well as the Middle-Bronze-Age (MBA) Babyne (Multi-Cordoned Ware, 2100-1600 BCE) archaeological groups (Table 1, S1 File). Five individuals came from the Dubynove kurgan, eight from the Katarzhyno kurgan 1, one from the Katarzhyno kurgan 2, five from the Liubasha kurgan, four from the Revova kurgan, and one each from the Mayaky kurgans 8 (M8.7) and 10 (M10.2.2).

**Table 1.**
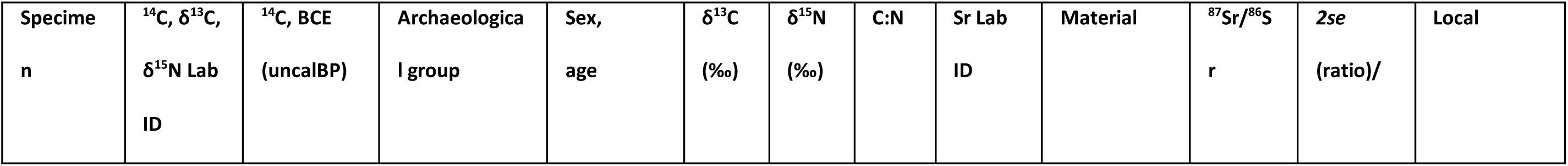
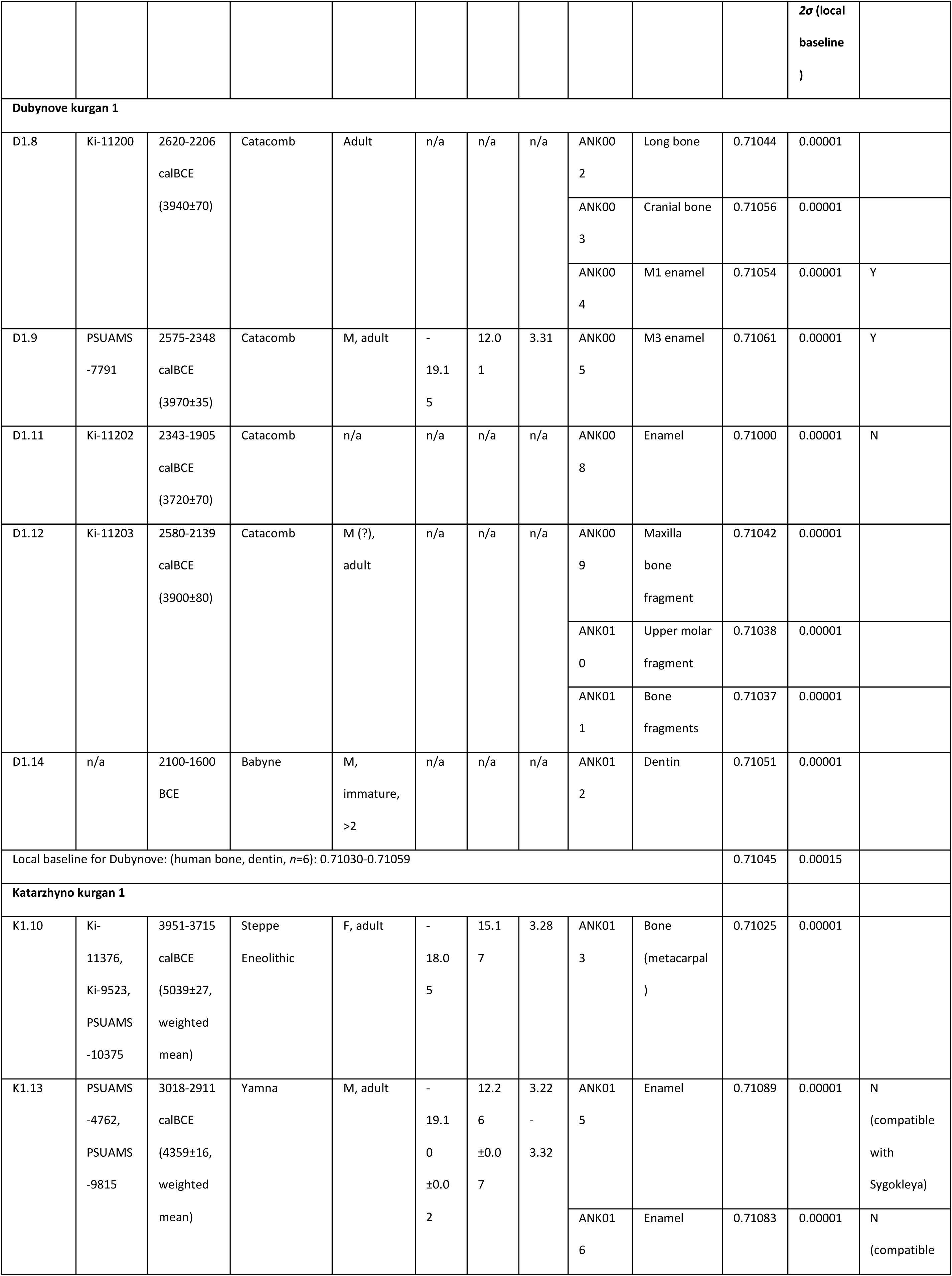
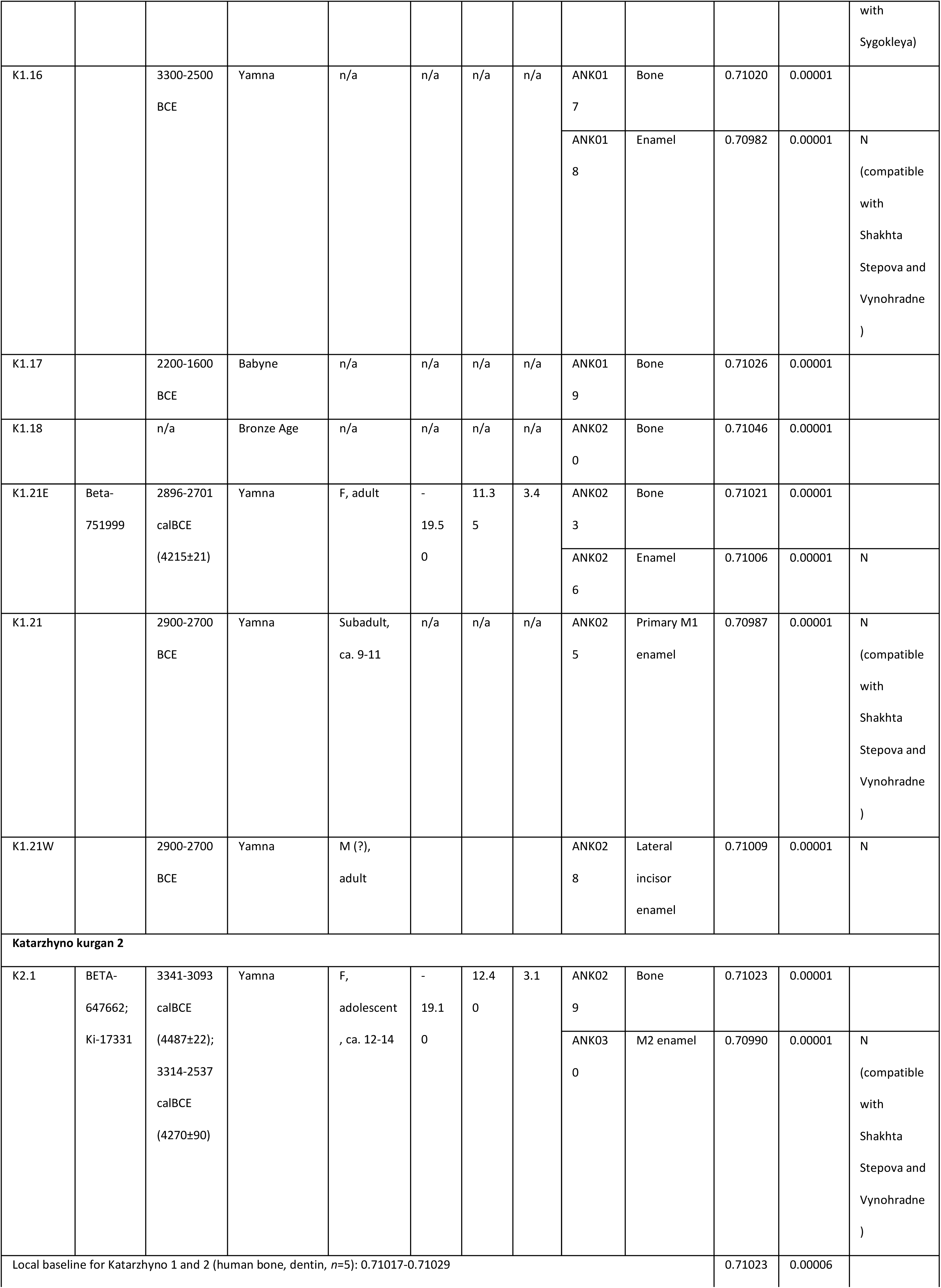
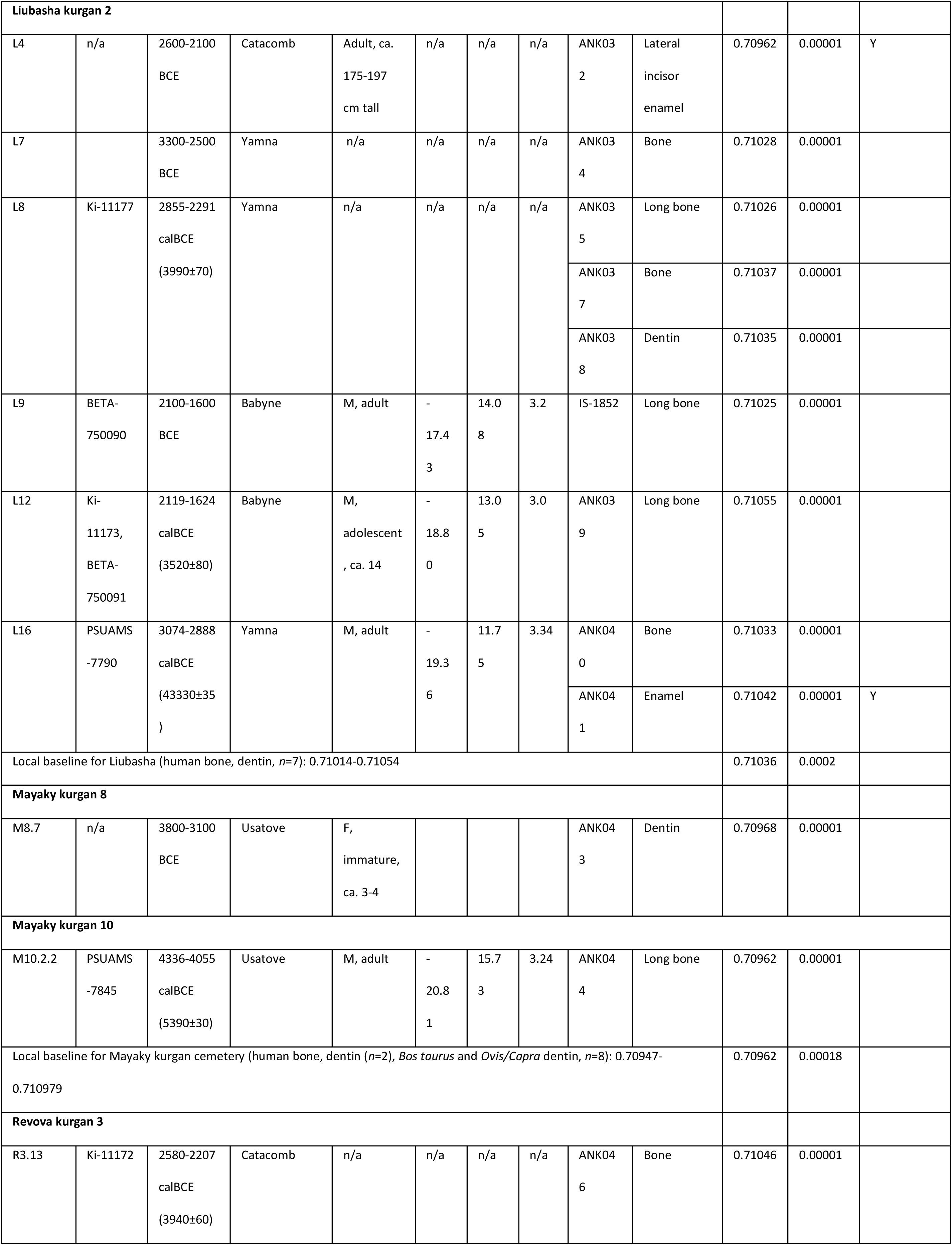
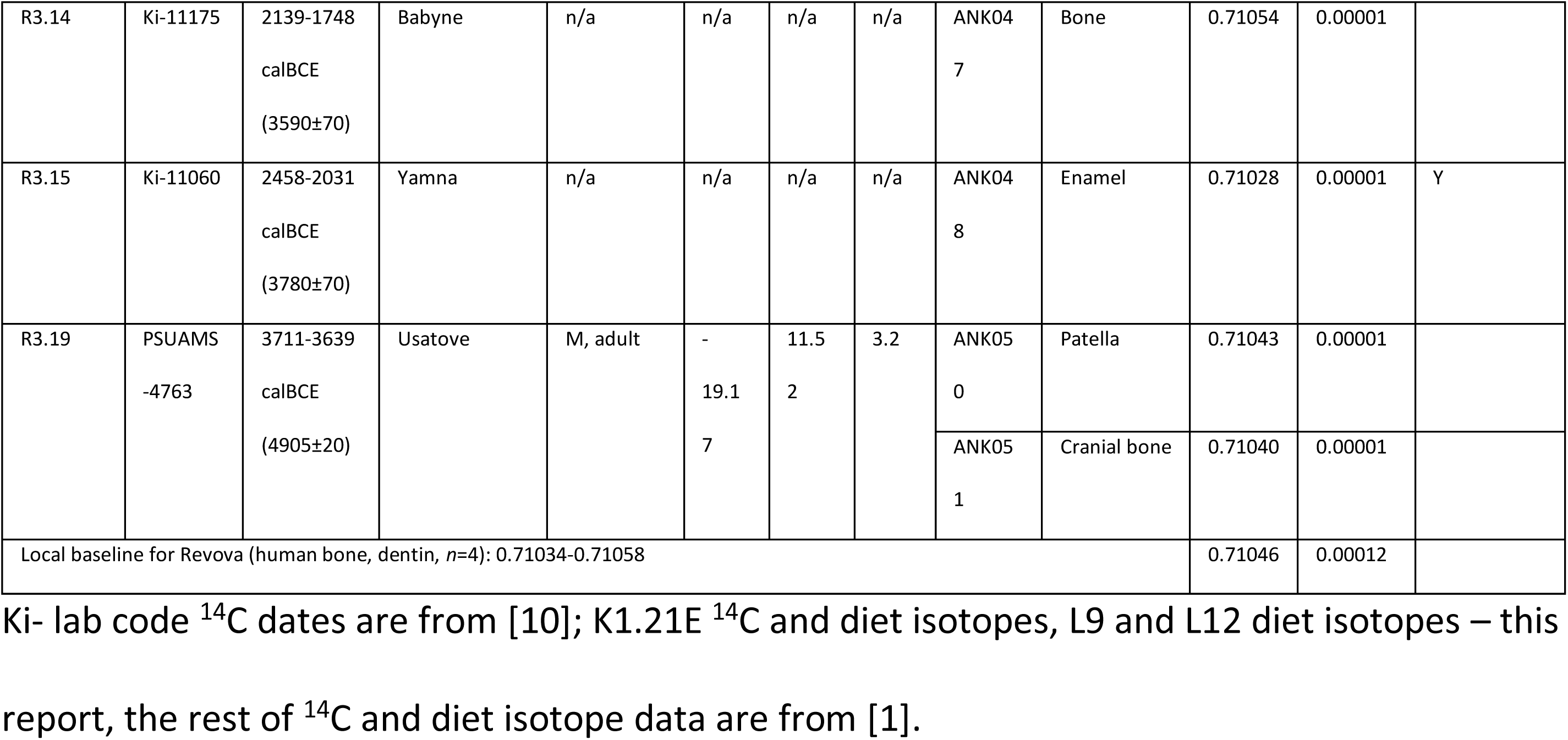
Strontium ^87^Sr/^86^Sr ratios produced in this study along with other available isotope metrics for 26 Eneolithic-Bronze Age individuals from the northwestern North Pontic Region.

### FTIR analysis (quality control)

For 12 out of 32 bone and tooth samples for which sufficient material was available, we assessed preservation of biogenic hydroxyapatite (bioapatite) using attenuated total reflectance Fourier transform infrared spectroscopy (ATR-FTIR) prior to Sr isotopic analysis to evaluate potential diagenetic alteration. Approximately 1 mg of powdered bone/tooth previously treated with 2% NaOCl and 0.2 N acetic acid from each sample was analyzed using a Thermo Scientific Niolet Summit iS5 FTIR spectrometer equipped with an Everest diamond ATR crystal. The spectra were collected in the mid-infrared range (4000-400 cm^-1^) at 4 cm^-1^ resolution, with 16 scans averaged per spectrum. We generated background spectra prior to the analytical session. All spectra were baseline-corrected before peak-height measurements. The peak heights were measured relative to defined baselines following established protocols for ATR-FTIR assessment of biogenic hydroxyapatite (e.g., [15,16]). Using the baseline corrected spectra, we calculated Crystallinity indexes (CI), also referred to as infrared splitting factor (IRSF), carbonate-to-phosphate ratios (C/P), and screened for potential calcite precipitation. All CI and C/P values were evaluated relative to established ATR-FTIR preservation ranges for archaeological bioapatite [15].

CI was used to evaluate crystal order and identify potential recrystallization of bioapatite. CI is calculated as:

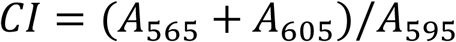

where A_565_ and A_605_ correspond to the heights of the phosphate V_4_ doublet peaks, and A_595_ corresponds to the height of the valley in between them. Elevated CI values can indicate a larger crystal size associated with diagenetic-related recrystallization. However, bone and tooth bioapatite CI values must be evaluated using separate criteria because of the naturally occurring differences in crystal size between these two tissues, which relate to their formation and their association with different proportions of organics. Our measured values were compared with published ATR-FTIR ranges for modern, well-preserved archaeological bioapatites.

The C/P ratios were calculated to assess carbonate incorporation and to detect potential exogenous carbonate contamination from the depositional environment. The ratio was determined using the formula:

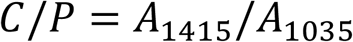

where A_1415_ corresponds to the B-type carbonate V_3_ band, and A_1035_ corresponds to the phosphate V_3_ band. Elevated C/P values can indicate the presence of exogenous carbonate.

All spectra were examined for the presence of secondary calcite formation which is identified by diagnostic absorption bands near ∼710 cm^-1^ and ∼874 cm^-1^. We recorded the presence or absence of calcite qualitatively.

Samples that exhibited CI and C/P values that were within published archaeological ranges and lacked calcite-associated absorption bands were considered suitable for subsequent mineral-based isotopic analyses.

### Strontium isotope analysis

⁸⁷Sr/⁸⁶Sr ratios for specimen L9 (Lab ID IS-1852) were produced by Isobar Science, Miami, FL, USA, following the methods in [17,18]. The rest of ⁸⁷Sr/⁸⁶Sr ratios was obtained at the University of Missouri. First, a chemical pretreatment was applied using the following sequence: soaking in a solution of 2% NaOCl, centrifugation, elimination of supernatant, rinsing with mQ water, soaking in 0.2N acetic acid, centrifugation, elimination of supernatant, rinsing with mQ water, drying. The samples were then placed in PFA vials and dissolved using 7M Optima Grade HNO_3_ at 110°C for 24 hours. The solutions were then evaporated at 90°C and the dry residues were re-dissolved in 2ml of 7M HNO_3_. The Sr extraction was conducted using Eichrom Sr-spec™ resin and 2 ml-reservoir columns following a protocol adapted from[19]. The Sr eluates were evaporated at 90°C. The dry residues were dissolved in 0.25 ml 14N HNO_3_ and evaporated at 90 °C. The residues were dissolved in 0.05N HNO_3_ before analysis on the multi collector - inductively coupled plasma-mass spectrometer in operation at the University of Missouri Research Reactor (Nu Plasma II, Nu Instruments).

The instrument was optimized daily to obtain a maximum intensity for ^88^Sr (∼ 7 V). The sample and standard solutions were prepared to obtain approximately 200 ng g-1 Sr. The isotopic standard SRM987 was measured multiple times at the beginning of each analytical session and after every two samples. The average and standard deviation for the SRM987 is 0.71023 ± 0.00004 (2SD) (n = 80). The values were corrected for mass fractionation using the 0.1194 value for the ^86^Sr/^88^Sr natural ratio. The isobaric interference at mass 87 was corrected using the 0.3857 value for the ^87^Rb/^85^Rb natural ratio. The isobaric interferences at masses 86 and 84 (^86^Kr and ^84^Kr) were corrected using the iterative method. The sample values were corrected by standard bracketing using the ^87^Sr/^86^Sr ratio value of 0.710248 [20]. To control the reproducibility of the entire process one duplicate (i.e., the entire procedure was applied twice to the same sample), one replicate (i.e., second analysis of the same solution) and two aliquots of the SRM1400 (Bone Ash) were analyzed. The values for SRM1400 were 0.71314 +- 0.00001 (2*se*) for each aliquot.

### Radiocarbon dating and stable isotope analysis

Newly reported ^14^C date for K1.21E as well as δ^13^C and δ^15^N isotope ratios for L9 and L12 were produced by BETA Radiocarbon lab, Miami, Fl following in-house protocols. The rest of δ¹³C and δ¹⁵N isotope data came from published datasets and from current study (Table 1, S1 Table).

δ¹³C and δ¹⁵N of bone collagen were analyzed descriptively and compared across regional groups using group means and standard deviations. No parametric tests were applied to δ¹³C and δ¹⁵N data due to sample size constraints. Instead, patterning was evaluated in relation to established regional isotopic ranges reported in the literature.

Statistical calculations were performed using online resources and Microsoft Excel for Mac Version 16. Graphs were constructed using Excel.

### Statistical analysis

Statistical calculations were performed using online resources (https://www.socscistatistics.com) and Microsoft Excel for Mac Version 16. Graphs were constructed using Microsoft Excel (Microsoft 365 for Mac, Microsoft Corp., Redmond, WA, USA). Descriptive statistics, including mean, standard deviation (SD), and range, were calculated for each isotopic group. Group-level summaries are reported as mean ±1 SD unless otherwise specified.

### Strontium baseline evaluation

According to [21], up to 100% of non-enamel strontium in buried remains can be diagenetic, thus representing the soil environment of the burial site. Rather than representing strictly local bioavailable strontium [22], and taking into account the results of the FTIR analysis, ratios derived from bone and dentine were interpreted as population-averaged values reflecting individuals’ movements across the broader region surrounding each kurgan during approximately the last 7–10 years of life, under broadly comparable dietary regimes.

Following [22], we estimated the baseline range for Dubynove, Katarzhyno 1/2, Liubasha, and Revova kurgans as the average of bone and dentin strontium ratios at each site ±2SD. Strontium ratios from specimen K1.18 were excluded from baseline calculations due to unclear archaeological context. We emphasize that the inland kurgan baseline represents a population-averaged mobility envelope rather than a strictly environmental bioavailable baseline. Because pastoralist groups likely circulated within a shared regional isotopic catchment, bone/dentin values are interpreted here as reflecting the isotopic space of habitual movement rather than a sedentary locality. The Mayaky kurgan cemetery baseline was estimated using the dentin values from M8.7 and the bone values from M10.2.2 as well as dentin from *Bos taurus* and *Ovis/Capra* [23], the latter reflecting local bioavailable strontium rather than population-averaged human mobility signatures.

Enamel values were evaluated relative to the obtained bone- and dentine-based averages, with concordance or divergence between tissues used to identify patterns of residential stability versus mobility or migration. Individuals were classified as “local” when enamel ^87^Sr/^86^Sr values fell within the population-averaged baseline range (±2SD) established for the kurgan or regional cluster under consideration (Table 1).

We compared the baseline and enamel ratios of strontium obtained from our sample selection to published strontium data from human bone, dentin and enamel from elsewhere in the Pontic steppe. Comparison sites included Pishchanka, Shakhta Stepova, Sygokleya, and Vynohradne from the east North Pontic steppe and the interred represented Steppe Eneolithic and EBA including Yamna and Catacomb archaeological groups) [25].

To assess whether the distribution of enamel classifications (within vs. outside baseline) differed significantly from dentin classifications, contingency tables were constructed and evaluated using chi-square (χ²) tests. Statistical significance was assessed at α = 0.05. Given the modest subgroup sample sizes, statistical results are interpreted cautiously and are used to identify structured patterning rather than to infer population-level parameters.

## Results

We used ATR-FTIR analysis to evaluate bioapatite preservation for 12 bone and tooth samples prior to conducting ⁸⁷Sr/⁸⁶Sr isotope analysis (Table 2). The CI values ranged from 3.25 to 4.46 (mean ± SD = 3.72 ± 0.35). The values of 10 samples all fall within the published ranges for both modern and well-preserved archaeological bioapatite and do not indicate extensive diagenetic recrystallization. The C/P ratios ranged from 0.00 to 0.33 (mean ± SD = 0.12 ± 0.09). All values were within the established ranges for archaeological bioapatite and do not suggest the presence of any exogenous carbonate material. No samples exhibited diagnostic absorption bands indicative of secondary calcite precipitation (∼710 cm^-1^ and ∼874 cm^-1^). Two of the samples had slightly elevated CI values. However, this likely reflects diagenetic crystal maturation rather than secondary carbonate contamination. Therefore, we have interpreted these as moderately recrystallized but chemically well-preserved, as further supported by the acceptable C/P ratios and the absence of calcite in the spectra. Overall, CI and C/P values, along with the absence of calcite-associated absorption bands, indicate minimal mineral diagenesis in the analyzed specimens. All samples were therefore considered preserved enough for subsequent ⁸⁷Sr/⁸⁶Sr analysis.

**Table 2.** Results of FTIR-ATR analysis.

| Sample Number | Peak 1415 cm<sup>-1</sup> | Peak 1035 cm<sup>-1</sup> | C/P Ratio | Calcite Peak at 710 cm<sup>-1</sup> | Peak 565 cm<sup>-1</sup> | Peak 605 cm<sup>-1</sup> | Peak 595 cm<sup>-1</sup> | CI Value |
| --- | --- | --- | --- | --- | --- | --- | --- | --- |
| ANK008 | 0.01 | 0.03 | 0.33 | No | 0.01 | 0.009 | 0.005 | 3.80 |
| ANK013 | 0.04 | 0.32 | 0.13 | No | 0.092 | 0.079 | 0.045 | 3.80 |
| ANK026 | 0.03 | 0.17 | 0.18 | No | 0.06 | 0.05 | 0.03 | 3.67 |
| ANK029 | 0.03 | 0.33 | 0.09 | No | 0.097 | 0.079 | 0.043 | **4.09\*** |
| ANK034 | 0.03 | 0.39 | 0.08 | No | 0.119 | 0.095 | 0.048 | **4.46\*** |
| ANK038 | 0.01 | 0.04 | 0.25 | No | 0.004 | 0.003 | 0.002 | 3.50 |
| ANK041 | 0.02 | 0.18 | 0.11 | No | 0.064 | 0.049 | 0.031 | 3.65 |
| ANK046 | 0.03 | 0.4 | 0.08 | No | 0.115 | 0.09 | 0.059 | 3.47 |
| ANK048 | 0.01 | 0.12 | 0.08 | No | 0.04 | 0.03 | 0.02 | 3.50 |
| ANK050 | 0.01 | 0.05 | 0.20 | No | 0.007 | 0.006 | 0.004 | 3.25 |
| ANK051 | 0.04 | 0.23 | 0.17 | No | 0.07 | 0.06 | 0.04 | 3.25 |
| ANK051_dupl | 0.02 | 0.12 | 0.17 | No | 0.037 | 0.033 | 0.019 | 3.68 |

\* Values slightly outside of acceptable range.

We generated 37 new ⁸⁷Sr/⁸⁶Sr ratios, ranging from 0.70962 to 0.71089, with an average of 0.71028±0.0003 (Table 1). Enamel samples D1.8, D1.9, L4, L16, and R3.15 of the thirteen used in the analysis fell within this population-averaged range for the western North Pontic region. Overall, the samples analyzed in this study fall within the 0.709–0.711 range previously reported for human skeletal tissues from the North Pontic region [22,25–27], indicating agreement with the currently available regional strontium isotope data.

Population-averaged baseline ranges based on bone and dentin were tightly constrained within each kurgan cluster (Table 1), supporting their use for evaluating enamel-derived mobility signals. No clear association between sex or age and mobility status was observed, although the limited sample size precludes robust demographic inference. Several enamel values fell within overlapping baseline ranges of multiple inland kurgans, reflecting the geological homogeneity of large portions of the northwestern steppe and underscoring the importance of regional rather than strictly site-specific interpretations.

Enamel strontium signature for the D1.8 and D1.9 Catacomb individuals from the Dubynove kurgan were in the local for that kurgan range. The strontium isotopic composition of the enamel from individual D1.8 was within the range defined for Dubynove and similar to that of the bone from the same individual. Enamel sample from burial D1.9 was at the upper end of the Dubynove baseline variation. We have interpreted this ratio to be within the local range, considering the uncertainty in the yet unsampled baseline range. A late Catacomb individual D1.11 was outside of the Dubynove kurgan baseline range and just outside of the upper end of the variation of the Shakhta Stepova kurgan from the eastern Pontic steppe [25] (Figs 1,2).

**Fig 2.**
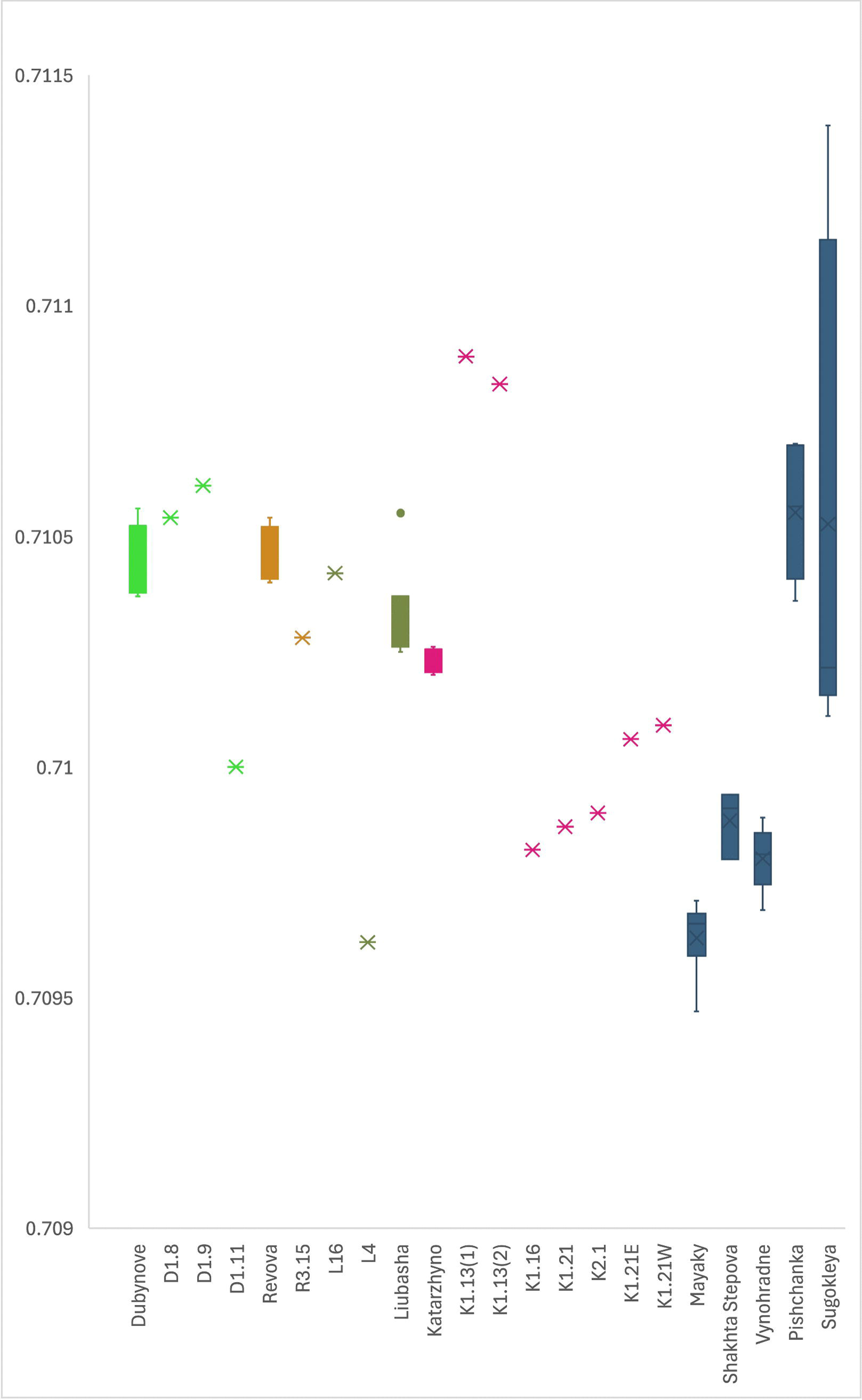
Enamel strontium ratios of individuals from western North Pontic steppe kurgans plotted against the baseline ranges of the North Pontic steppe established on bone and dentin. The northwestern steppe baseline is represented by Dubynove, Revova, Liubasha, Katarzhyno, and Mayaky (this report). The eastern steppe baseline is represented by Shakhta Stepova, Vynohradne, Pishchanka, and Sugokleya [25].

Enamel sample from the Yamna R3.15 individual differed from the Revova kurgan range but fell within the range of the Liubasha and Katarzhyno kurgans baseline. Enamel from the Liubasha primary burial L16 was within the Liubasha kurgan range as well as the ranges for Dubynove and Revova, while the enamel from the Liubasha’s Catacomb burial L4 was below this range but fell within that of the Mayaky Eneolithic archaeological site (Fig 2).

None of the enamel from the Katarzhyno 1 and 2 kurgan samples matched the range of that kurgan group or that of the northwestern Pontic baseline established by the bone and dentin strontium ratio ranges from the six kurgans from this report (Fig 2). The three Yamna adult individuals (K1.16, K1.21E and K2.1) for whom both enamel and bone or dentin were measured show a discrepancy between the tissues, which is indicative of migration between the moment the enamel formed and the last 7-10 years of life. Except for K1.13, enamel samples from burials at Katarzhyno kurgans were within the baseline ranges for the eastern North Pontic steppe [25]. Enamel from Yamna individuals K1.16 adult and the K1.21 subadult as well as the adolescent early Yamna K2.1 from the primary burial at the Katarzhyno 2 kurgan were within the baseline variation of Shakhta Stepova and Vynohradne. Enamel strontium ratios of the Yamna K1.21E and K1.21W adults did not overlap with any baseline range in Fig 2. Enamel of individual K1.13 fell within the range of Sugokleya.

To explore whether stable dietary isotopes might provide additional insights into mobility and possible affiliations with kurgan populations from the greater steppe region, we compared the δ¹³C and δ¹⁵N values obtained in this study with published data for Eneolithic–Bronze Age individuals from the eastern North Pontic steppe as well as the West Caspian steppe [25,28–30]. Eneolithic-EBA δ^13^C values from northwest Pontic steppe were significantly different from the eastern Pontic steppe, while δ^15^N values were not (Mann-Whitney Two-Tailed test z-scores = 3.45944/ -0.45691, *p* = 0.00054/0.64552, respectively). We thus plotted the means for these values separately on the graph Fig 3. As evident from the graph, and confirmed statistically, all but two Eneolithic-Bronze Age samples from our sample selection were within the western North Pontic steppe distribution.

**Fig 3.**
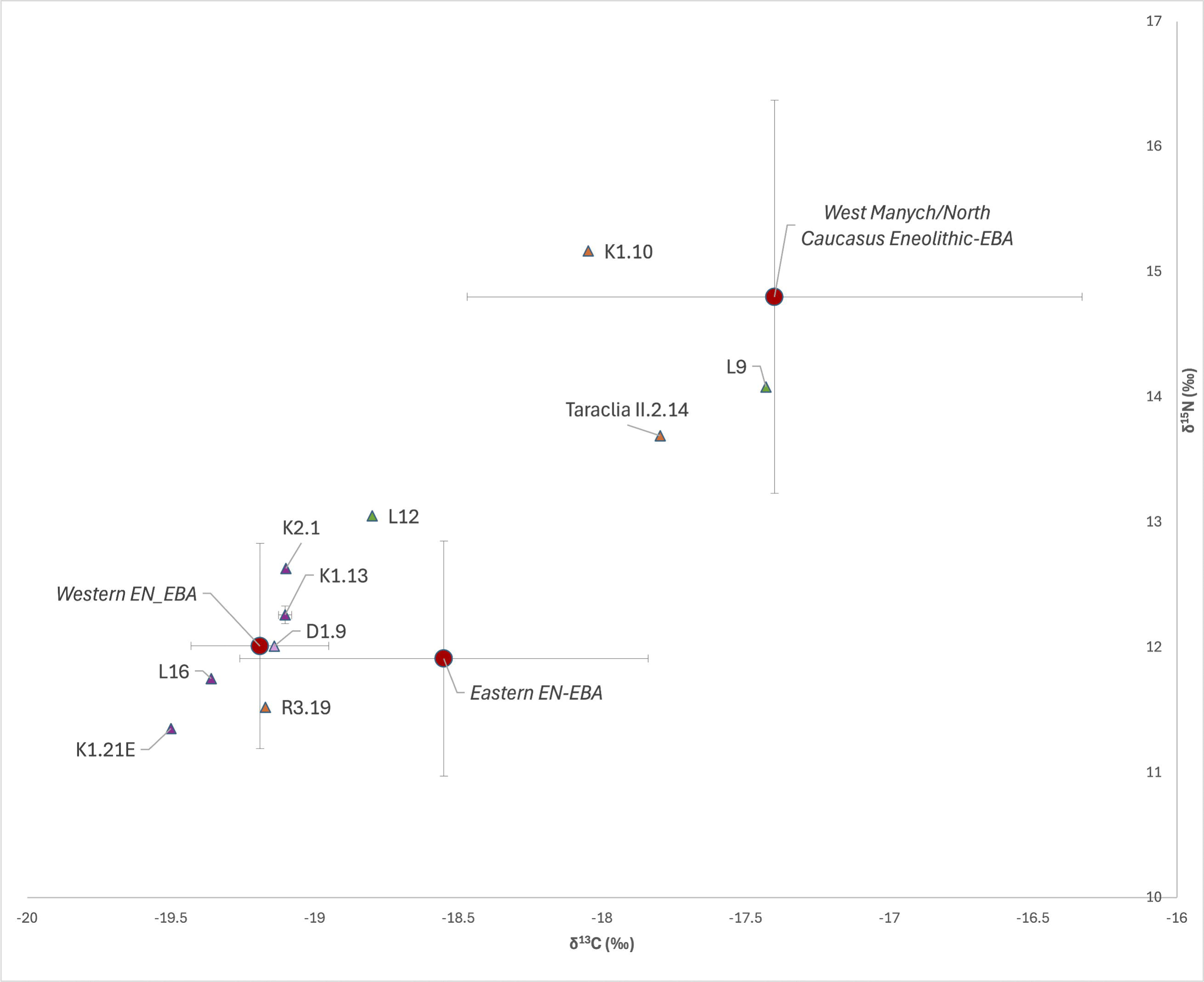
Diet isotope ratios of Eneolithic-Bronze Age populations of the North Pontic and West Caspian-North Caucasus steppe. Isotope values used to construct the graph, and the corresponding references are listed in S1 Table of S1 File.

Of the two identical twins L9 and L12 from the Babyne archaeological group of the MBA, the isotopic composition of individual L12 falls well within the variation of Eneolithic-EBA populations of the eastern Pontic (joint χ²(2) = 1.60, p = 0.45) and was also compatible with those of the northwest Pontic steppe (joint χ²(2) = 4.25, p = 0.12). On the other hand, the isotopic composition of individual L9 situated well outside the variation of both comparison groups (χ²(2) = 7.82, p = 0.02, and χ²(2) = 60.15, p < 0.001, respectively), indicating a diet distinct from either reference population. At the same time, L9’s δ^13^C and δ^15^N values lie within the isotopic ranges of both West Manych Yamna and Eneolithic steppe Maykop, with joint probabilities of p = 0.72 and p = 0.96 respectively, suggesting strong isotopic similarity to these west Caspian/ North Caucasus Piedmont Eneolithic-Bronze Age groups.

Steppe Eneolithic individual K1.10 from the primary burial at the Katarzhyno 1 kurgan was another outlier in terms of δ^13^C and δ^15^N values, showing a similar to the L9 trend. The stable isotope values of K1.10 (δ¹³C = −18.05‰; δ¹⁵N = 15.16‰) are statistically incongruent with Eneolithic-EBA populations of the eastern and western Pontic steppe (χ²(2) = 12.49, p = 0.002, and χ²(2) = 37.43, p ≪ 0.001, respectively), but fall within the isotopic ranges of both West Manych Yamna and Eneolithic Caspian steppe (χ²(2) = 0.52, p = 0.77, and χ²(2) = 0.59, p = 0.75), suggesting affinity with the latter groups.

## Discussion

Kurgan burial tradition began in the North Pontic steppe in the early 4^th^ millennium BCE [1,31]. Kurgans were re-used for hundreds of years thereafter. Most of the Eneolithic-EBA kurgans in the North Pontic steppe contain multiple individual burials. The western North Pontic kurgans studied in this report contained 5-21 burials of mostly single individuals. One of the burials in the Katarzhyno kurgan 1 contained two adults and a subadult, being one of the only two such burials in the northwest Pontic steppe among the 2000 surveyed intact burials [32]. It is unknown at this point if the subadult is genetically related to the adults in the same burial. Some kurgans functioned as burial places into the modern times [10].

Bone and dentin strontium values from the Bronze Age individuals interred in the five inland kurgans in the present sample selection were mostly concordant and within a relatively tight isotopic baseline range of 0.71017-0.71059 (Table 1). In contrast, their enamel strontium ratios largely deviated from this baseline, suggesting childhood residence in geological environments distinct from those represented during the final years of life. Yamna individuals from the Katarzhyno kurgan group, which provided the bulk of enamel datapoints for this study, show distinctive trends in the enamel strontium signature variation. Such as individual K1.13 was isotopically distinct from the northwest Pontic baseline and the remaining Katarzhyno individuals, with a strontium signature within the baseline range of the Sugokleya kurgan in the eastern steppe. Individuals K1.16 and K2.1 show parallel shifts between enamel and bone values, with closely matching enamel ratios that point to a shared area of origin outside of the northwest Pontic. The K1.21 subadult displayed a similar enamel signature, suggesting that the three grew up in an area geologically similar to that of the Pishchanka and Vynohradne kurgans in the eastern steppe. The intermediate enamel values observed in individuals K1.21E and K1.21W are best explained by mobility during enamel formation, resulting in isotopic signatures that average residence across two geologically distinct regions. The variability of enamel signatures from Katarzhyno may reflect seasonal mobility of the pastoralist Yamna groups. At the same time, if they all followed the same mobility pattern, one would expect that average the same regions at the same time, resulting in a similar isotopic signature. Alternatively, the interred in the Katarzhyno kurgans may have originated from different steppe regions with distinct mobility patterns, and comingling in the Katarzhyno kurgans area.

In contrast, the enamel strontium signature of Yamna individual L16 from the primary burial in the Liubasha kurgan suggest L16 to have been part of the locally established Yamna group. Enamel strontium ratios from the Liubasha’s burial L4 of the succeeding Catacomb archaeological culture suggests L4 spent the middle childhood in an area geologically similar to the coastal area around the Mayaky site. The analysis of archival excavation reports and pertinent publications revealed a number of archaeological sites associated with the Catacomb culture group along the Dniester Estuary and the adjacent Black Sea littoral. Thus, it is possible that L4 may have been associated with a Catacomb population group that utilized these sites.

Strontium baseline for the inland kurgans in our study was overlapping, although the ranges were somewhat dissimilar in breadth, likely due to limited number of baseline samples from each kurgan. There was an overlap between the inland kurgan group in our selection and the lower isotopic strontium ratio range of the Pishchanka kurgan in eastern Ukraine (Fig 2), possibly due to similar geology, which around Pishchanka consists of Cenozoic sediments and exposed Proterozoic plutonic rocks [25]. The area around the Sugokleya kurgan encompassed most of the known bioavailable strontium variation in Ukraine, likely explained by having a complex geology (Fig 1), including a unique combination of crystalline Precambrian and sedimentary layers, tectonic features, igneous intrusions, and ore formation [35–37]. Adding to the uniqueness of the area’s geology is its association with the 24km Boltysh crater, thought to had been formed by an asteroid that followed the Chicxulub impact at the Cretaceous-Paleogene boundary ca. 66 MYA [38].

δ^13^C and δ^15^N values of all but two of the studied individuals from the inland kurgans were within the dietary range of the Eneolithic-Bronze Age inhabitants of the North Pontic steppe (Fig 3). δ^13^C and δ^15^N values of Eneolithic individual K1.10 and the MBA Babyne individual L9 indicate that they spent the last part of their lives outside of the North Pontic, possibly in the west Caspian area. Supporting a west Caspian dietary affiliation for some mobile steppe individuals from the North Pontic, the Eneolithic Taraclia II.2.14 individual interred in the western steppe, identified as a second cousin of Sharakhalsun burial 18 from the Manych steppe [1], exhibited δ^13^C and δ^15^N values consistent with the west Caspian steppe range (Fig 3).

At the same time, δ^13^C and δ^15^N values of L12, the L9’s identical twin, are within the North Pontic steppe range. This suggests that L12, an adolescent, likely lived and died in the North Pontic, while L9, who lived into adulthood, may have spent the last decade of life in the North Caucasus-West Caspian area, but was buried near his likely birthplace in the western steppe in the same kurgan as his twin brother. On the other hand, L9 could have sustained on a special diet. Alternatively, L9’s δ^13^C and δ^15^N values may reflect population-level dietary changes, possibly associated with the North Atlantic cooling event at c. 2250 BCE/4200 BP [39], followed by a rapid climate change c. 2250-1850 BCE (4200-3800 BP [40]).

Comparative diet isotope information for the Babyne archaeological group is limited. Anthropological studies indicate that Babyne individuals experienced considerably higher levels of physiological stress compared to steppe populations from the preceding chronological period [41]. Nutritional stress has been shown to lead to elevated δ¹⁵N ratios in humans [42] as seen in L9, but the pattern is less consistent and smaller in magnitude for δ¹^3^C. If climate change were the cause of shifts in δ^13^C and δ^15^N values, then these data could be used as a correlate of environmental dynamism. The limited data available on the Babyne diet and the scarcity of climatic data for the North Black Sea region during the MBA hinder a detailed evaluation of the latter hypothesis.

Using data obtained in this report in combination with those available from literature, we constructed first strontium baseline heatmap for the North Pontic region during the Eneolithic-MBA chronological period (Fig 4). Additional analyses of strontium isotope ratios across the North Pontic region, particularly through denser spatial sampling of both human remains and environmental proxies, will expand this baseline framework, improve discrimination between overlapping geological zones, and allow more precise reconstructions of mobility patterns, seasonal movement, and population connectivity within the steppe landscape that played a central role in continental-scale cultural and demographic transformations. As a pivotal corridor linking the Carpathian Basin, the Balkans, the Caucasus, and the Volga-Ural region, the North Pontic steppe served as a dynamic center where populations interacted, merged, and reorganized. Expanding strontium-based provenance studies here would therefore clarify not only patterns of movement, but also the processes through which this region became a major arena of change in prehistoric Eurasia.

**Fig 4.**
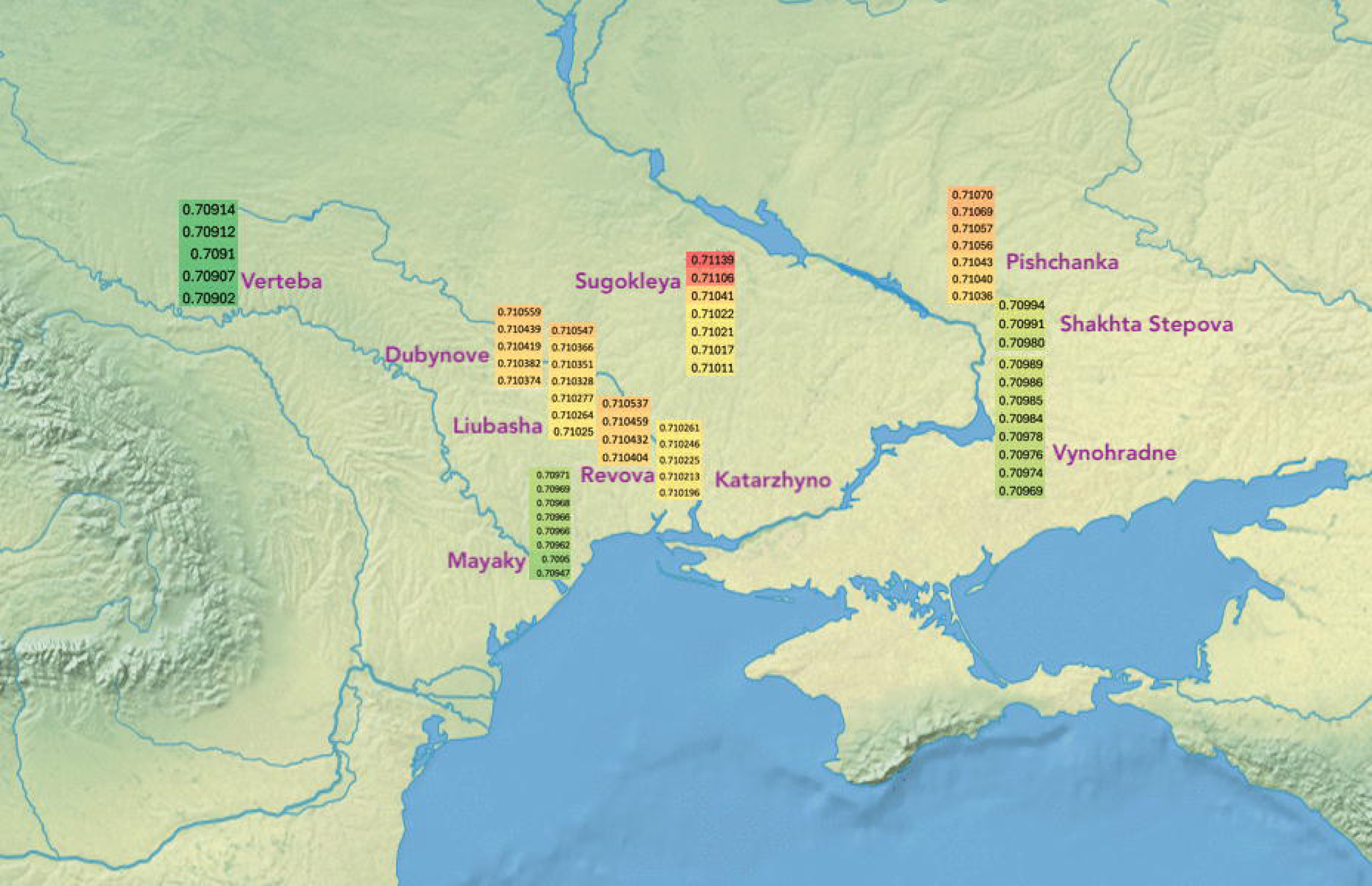
A heatmap of ^87^Sr/^86^Sr baseline distribution in the North Pontic region during c. 4000-2000 BCE chronological period based on the current study and the data available from literature. Verteba baseline represents values from water and snail shells from a ritual site of Eneolithic Trypillia culture [43]. Values for Dubynove, Katarzhyno, Liubasha, Revova (this report), Pishchanka, Shakhta Stepova, Sugokleya, and Vynohradne [25] are from human bone and dentin. Values for Mayaky are from human bone and dentin as well as *Bos* and *Ovis/Capra* dentin [23].

## Acknowledgements

The authors thank the Odesa Archaeological Museum for providing access to the samples studied in this report.

## Supporting Information

S1 File. Description of sites and burials.

S1 Table. Stable isotopes of carbon and nitrogen used to construct the graph in Fig 3.

