## Supplementary File for "Tracing mobility among Eneolithic-Bronze Age Kurgan populations in the North Pontic steppe"

### S1 File. Description of sites and burials

The kurgans sampled in this study were situated in the Odesa Oblast of Ukraine, on a roughly 190-kilometer north-south line stretching from the Dniester Estuary. (Fig 1). The Dubynove kurgan was located in the southern part of the forest-steppe zone, the rest were in the steppe zone. Detailed descriptions of kurgans and excavations are provided in [1–4]. With the exception of the Mayaky mounds, these kurgans were used for burials over periods exceeding 1000 years.

#### Dubynove Kurgan 1 (48.133, 30.282)

Podilskiy District

Kurgan 1 was located on the plateau of the west bank of the Pivdennyi Buh River, 200 m east of the village of Dubynove. The embankment was disturbed by earthworks, the top was cut off, the slopes were repeatedly plowed. Around the kurgan, an extensive hollow was observed at the place of the ancient excavation of the bulk soil. From the level of this depression, the height of the mound was 2.0–2.2 m, with a diameter of 36 m. The kurgan contained 15 burials. Burials 8, 9, 11, 12 (Catacomb archaeological culture) and burial 14 (Babyne archaeological group) are presented in this report.

Based on the available $^{14}C$ dates, the Dubynove Kurgan 1 was in use by the Catacomb archaeological culture for 52±14 years. Considering the undated Eneolithic primary and Late Bronze Age burials, the kurgan's use may have extended over 1700 years.

##### Dubynove Kurgan 1 burial 8 (D1.8)

Adult, 2620-2206 calBCE (3940±70BP, Ki-11200) [2]

Burial 8 was found 9 m to the south-southeast of the benchmark, at a depth of 2.34 m. The 2.25 × 1.75 m catacomb burial chamber oriented along its long axis in the southwest-northeast direction. The vault at the entrance to the chamber, presumably, reached a height of 0.75 m. The entrance well was apparently located on the south side of the chamber, where the walls were made strictly vertical. The interred was laid extended on the back, head to south-west. The hands were slightly bent at the elbows and placed along the body; the left hand was on the pelvis. A grayish-brown pot of squat proportions, with a short neck, a slightly curved rounded rim, pronounced shoulders and rounded sides tapering to a flat bottom was found behind the head, under the wall of the chamber.

##### Dubynove Kurgan 1 burial 9 (D1.9)

Male, adult, 2575-2348 calBCE (3970±35 BP, PSUAMS-7791) [1]

Burial 9 was found 8 m to the southeast of the benchmark, at a depth of 2.55 m. The long side of the 2.1 × 1.4 oval burial chamber was oriented to the south-southwest – north-northeast. The entrance well was apparently located at the eastern side of the chamber. The interred was laid extended supine, with the head to the south-southwest, face turned to the east. The left wrist was placed on the pelvis. There was no inventory.

##### Dubynove Kurgan 1 burial 11 (D1.11)

2343-1905 calBCE, 3720±70 BP (Ki-11202) [2]

Burial 11 was found 14 m to the southwest of the benchmark, at a depth of 2.7 m. The trapezoidal 1.5 × 0.75–1 m burial chamber oriented in the east-west direction was a vaguely traced catacomb or a pit. The walls of the chamber have been preserved to a height of 0.6 m (total depth of 3.3 m). The interred was lying on the back, almost diagonally across the camera, with the skull to the northwest.  The legs, initially raised with the knees up, fell to the right. The burial position is characteristic of the Yamna culture. There was a deposit of pink ocher on the bones, and brown decay from an underlay at the bottom. In the south-eastern corner of the chamber stood an "amphora" with a straight neck, a spherical body and a flat bottom separated by a border. Two pairs of arched handles with vertical holes were symmetrically affixed to the shoulders. The edge of the neck was covered with notches.

##### Dubynove Kurgan 1 burial 12 (D1.12)

Male, adult, 2580-2139 calBCE (3900±80 BP, Ki-11203) [2,5]

Burial 12 was found 10 m south of the benchmark. Two processed stone blocks covering a 0.65 × 1 m entrance well on the eastern side of the catacomb, were discovered at a depth of 2.75 m. Before entering the burial chamber, the entrance well had a step 0.15 m high, covered with stone blocks. The 2.4 × 1.8 m burial chamber had the shape of an oval expanding to the north. The bottom of the chamber was 0.55 m below the bottom of the entrance well. The interred was lying on the back, slightly contracted to the left, head to the north, facing east, toward the entrance to the burial chamber. The legs were bent at the knees to the left. The left arm was extended along the body, the right arm was bent at the elbow, and the hand was lying in the pelvis. Brown rust is noted under and around the skull. Opposite the front part of the skull was a mace 5.6 cm high, 6.2–

6.9 cm in diameter with a hole for a handle attachment, made of greenish-black gneiss. The surface of the mace was polished and covered with red-brown ocher and chalk.

##### Dubynove Kurgan 1 burial 14 (D1.14)

Male, immature (>2), 2100-1600 BCE

Burial 14 was found 6.4 m to the east-southeast of the benchmark, at a depth of 2.8 m. The burial pit was not traced. The incompletely preserved bones of the skeleton indicated the interred lay contracted on the right side, head to the south, facing east. The legs were bent (the hip angle was obtuse), the right arm was extended with the wrist to the knees, the bones of the left were not preserved. Spots of red ocher were noted on the skull.

#### Katarzhyno kurgan group (47.025, 30.288)

Znamianka, Ivanovskyi District

The Katarzhyno kurgan group was located 4 km south of the village Znamianka (until 1932 - Katarzhyno), on the plateau of the west bank of the Malyj Kuyalnyk River. The group consisted of five kurgans. Four of them, including Kurgan 2, were arranged in a row from north to south. Kurgan 1, the largest of the five, was located a little off to the side. The Katarzhyno kurgan group was located on the northern end of a kurgan chain that stretched along the meridional watershed up to 120 m above the river thalweg.

##### Katarzhyno 1 Kurgan 1

Kurgan 1 had a diameter of 80–85 m, its height at the beginning of the excavations was 6.45 m, while according to the topographical survey in 1950, the height was 7.3 m. The slopes of the kurgan were heavily plowed and partially cut. The top was also cut off and disturbed by modern digging. A total of 21 burials were uncovered in the kurgan. Burials 10 (Endolithic, main burial); 13, 16, 21 (Yamna); 17 (Babyne); 18 (Bronze Age) are presented in this report.

The chronological span of use of the Katarzhyno Kurgan 1 based on the available $^{14}C$ dates between main Eneolithic burial K1.10 and chronologically youngest Yamna burial K1.1 is 1318±11 years. This does not take into account the undated Babyne burials within the kurgan, which would extend the kurgan's use another 230 to 880 years.

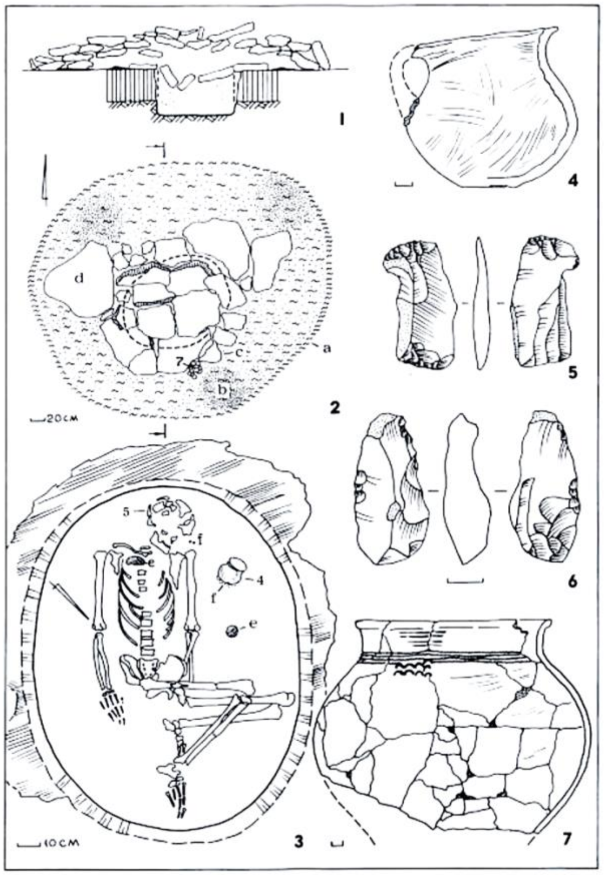

*Figure S1. Katarzhyno kurgan 1 burial 10. Image from[2], reproduced with permission.*

##### Katarzhyno Kurgan 1 burial 10 (K1.10) Female, adult, 3955-3783 calBCE (5039±27 BP) [1]

Burial 10 was discovered 6.5 m southeast of the central benchmark, buried beneath an embankment composed of several layers of earth and a layer of stone. At the level of the ancient day surface, the burial pit was covered with a reed mat containing preserved directions of the fibers and a single layer of limestone. Among the stone slabs, a fragment of an anthropomorphic stelae with a chiseled shape resembling a human head was found. The reed mat near the stelae was covered with a layer of 5-8 mm of organic matter in the form of a dense yellowish-white streak (possibly remnants of honey), which was further covered by a large fragment of a ceramic pot. Two other slabs without signs of processing were laid on the east side of the stone arrangement. The bottom of the tomb was covered with a dark brown decay of the bedding, on which the deceased was laid contracted on the back, with the legs were bent and the knees turned to the right. The arms were straightened and laid along the body with the palms down. The skull was tilted to the left shoulder, and the orientation was southwestern. A thin deposit of ocher was detected on the bones. A 2.5 × 6 cm lump of ocher was found near the left hand at the level of the elbow. Another 2.5 × 10 cm lump of ocher was found on a layer of coal placed on the chest near the right clavicle. The burial inventory consisted of a molded asymmetric askos-type vessel with an oval-shaped broken-off handle, a coarsely molded shell-tempered pot with a missing bottom and a corded ornament in the form of a horizontal four-row belt and bows attached from below, resting on a layer of coal close to the left shoulder, and a chisel-type tool with two working sides made of gray flint, found under the skull. The same type of tool with one side showing signs of use found in the grave fill.

##### Katarzhyno Kurgan 1 burial 13 (K1.13)

##### Male, adult, 3024-2908 calBCE (4359±16 BP)[1]

Burial 13 was discovered 5.6 m south of the benchmark, at a depth of 2.35 m. It was not possible to determine whether there had been a ledge in the burial chamber. The chamber was covered by four large unaltered limestone slabs. The chamber was rectangular in shape with rounded corners, measuring 2.0 × 0.9 m, and was oriented lengthwise from northeast to southwest. The depth from the covering level was 0.6 m (total depth: 2.95 m). The northern side of the pit was heavily damaged by rodents. The skeleton was poorly preserved: the right humerus was missing, the skull was shifted to the side, and the cranial vault was destroyed. The deceased was laid on the back, with the head to the northeast. The arms were extended along the body; the bent legs were positioned with the knees facing upward. At the bottom was a lining made of bark, sprinkled with chalk. An ocher deposit was found near the left shoulder.

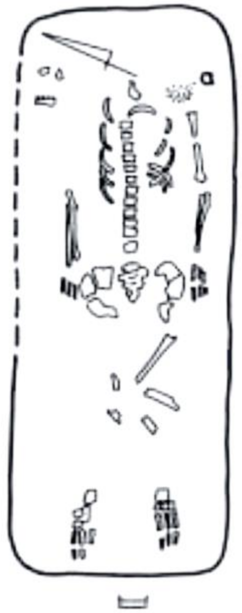

*Figure S2. Katarzhyno kurgan 1 burial 13. Image from[2], reproduced with permission.*

##### Katarzhyno Kurgan 1 burial 16 (K1.16)

##### Male, adult, 3300-2500 BCE

The rectangular pit of burial 16, measuring 1.65 × 1 m and with rounded edges, was discovered 30 m north of the benchmark at a depth of 5.1 m. The pit was covered with wooden planks placed lengthwise that sank into the burial chamber, with remains lying directly on the bones of the skeleton. The interred was laid on the back, head to the west. Arms were stretched along the body; legs knees bent to the right. A 5.2 × 2.5 × 0.7 cm blade made from light gray flint with white inclusions was found between the tibia and femur. Fragments of the bottom of a molded black vessel with admixtures of sand chamotte with grooved surface leveled with a spatula and containing remnants of brown organic material with an imprint of a fruit pit was found by the legs.

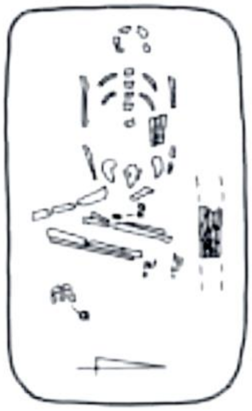

*Figure S3. Katarzhyno kurgan 1 burial 16. Image from[2], reproduced with permission.*

##### Katarzhyno Kurgan 1 burial 17 (K1.17)

##### Male, 2200-1600 BCE

Burial 17 was found 31.5 m south of the benchmark, at a depth of 5.7 m. The oval 1.4 x 1 meter burial chamber was traced to a depth of 0.2 m (total depth: 5.9 m), oriented east-west. The interred was laid on the left side, with bent legs, head to the east. The skeleton was poorly preserved, the bones of the arms are missing, the bones of the chest were displaced to the northern wall of the pit. In the place where they were originally supposed to be, the bottom of the grave was pierced to a depth of up to 3 cm in a section with a diameter of 0.25 m. The same burn was noted in front of the skull. The bones and bottom of the pit are covered with a layer of charcoal up to 5 cm thick. Several ribs were burnt.

##### Katarzhyno Kurgan 1 burial 18 (K1.18)

Burial 18 was discovered 19 meters southeast of the benchmark, at a depth of 3.4 meters. The contours of the pit were not traced. The skeleton was severely damaged by rodents. Based on the preserved remains, it appears that the deceased was laid with the head facing northeast and the legs bent at the knees and falling to the left. The position of the hands could not be determined.

##### Katarzhyno Kurgan 1 burial 21

##### 21E, adult, female (K1.21E)

##### 21W, adult (K1.21W)

##### 21, subadult (ca. 9-11) (K1.21)

The triple burial 21 was discovered 20 meters southeast of the benchmark, at a depth of 4.25 meters. The burial pit, oval in plan and funnel-shaped in cross-section, measured 1.8–2.0 meters in diameter at the top and 1.8–1.2 meters at the bottom. The pit, 1.3 meters deep from the level of the log ceiling (total depth: 5.55 meters), was covered with logs laid lengthwise, each about 0.2 meters in diameter, which collapsed into the pit. Fragments of logs were found both in the fill and on the bones of the skeletons. The burial contained two adults and a subadult. The adults lay in a flexed position on their backs, with their heads to the southwest. The eastern skeleton of a female (21E) was found with the arms extended along its body and the legs bent with the knees raised. Copper oxide traces were discovered on two of its ribs. The western skeleton (21W) had

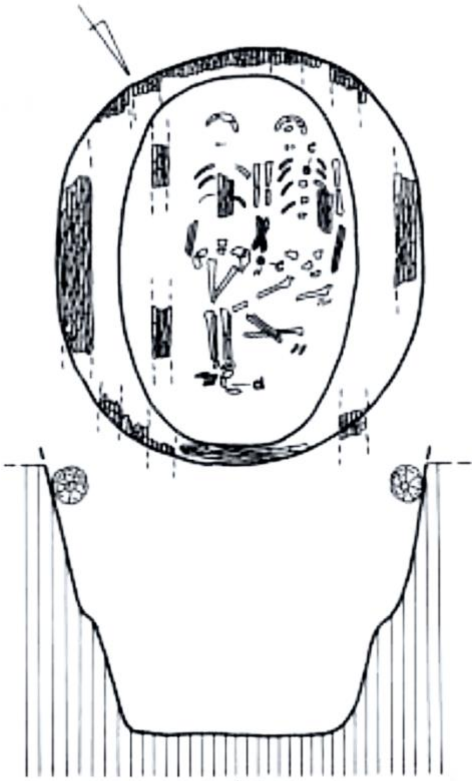

Figure S4. Katarzhyno Kurgan 1 burial 21. Image from [2], reproduced with permission.

the legs bent at the knees to the right. The left arm of skeleton 21W extended along the body, with the hand tucked under the pelvis. The right hand of skeleton 21W was placed on the left arm of skeleton 21E. At the wrists of the crossed arms, there was a lump of purple ocher, egg-shaped, about 4 cm in diameter. A 2 × 3 cm stick of the same ocher was found on the ribcage of skeleton 21W. On the feet of skeleton 21E lay the skull of a ca. 9-11-year-old subadult (skeleton 21). The subadult's postcranial skeleton was represented by a few fragments of ribs and long bones.

#### Katarzhyno Kurgan 2

Kurgan 2, 2.4 × 50 m, was located 20 m to the northwest of Kurgan 1. The embankment was heavily plowed, and its top was cut off. Three construction horizons were traced in the kurgan. The kurgan contained six burials.

##### Katarzhyno Kurgan 2 burial 1 (K2.1)

##### Female, adolescent (ca. 12-14), 3348-3038 calBCE (4490±30 BP, BETA-647662)

The primary burial 1 was discovered 1.4 m south of the benchmark, at a depth of 2.3 m. At this level, a layer of small flat stones (measuring 0.2–0.3 m) was recorded, arranged in one to two tiers and not extending beyond the boundaries of the grave. Beneath the stones, traces of a decayed wooden covering were observed. The burial chamber was rectangular in plan, measuring 1.7 × 1.1 m, with rounded corners and a depth of 0.6 m from the level of the ancient ground surface (total depth of 3.0 m). The individual was laid

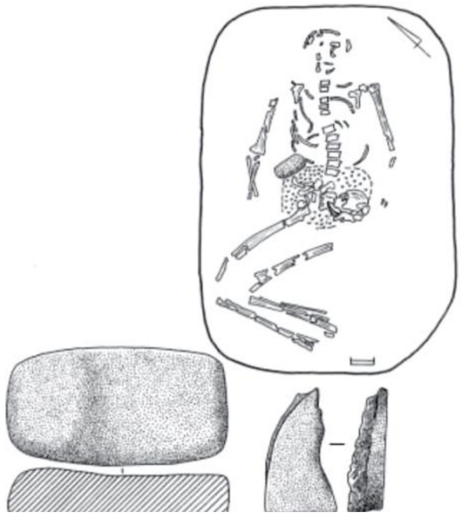

Figure S5. Katarzhyno kurgan 2 burial 1. Image from[2], reproduced with permission.

on the back, head oriented to the northeast. The arms were slightly spread to the sides, bent at the elbows and extended along the body, while the legs were bent at the knees and turned to the right. A gray sandstone slab of prismatic shape, measuring 11.7 × 6.3 × 3.4 cm was found between the right side of the rib cage and the pelvis. The tool had an upper surface polished from use; the bottom surface and all lateral edges were evenly covered with characteristic pitting. A semi-segmental 6.6 × 3.6 × 2 cm fragment of limestone slab with sharp edges (one of the edges notched) was found on the sacrum. A 1 cm-thick layer of green clay coating was detected below the pelvic bones and at the level of the lumbar vertebrae.

#### Kurgan Liubasha (47.557, 30.3143)

Novohryhorivka, Mykolaivskyi District

Kurgan Liubasha was located on the plateau of the west bank of the Tiligul River 118 m above the river thalweg. The kurgan stood at the edge of the highest watershed plane in the immediate vicinity, stretching from the southeast to the northwest between Tiligul and the Glubokiy Yar ravine. Other kurgans on the same section of the plateau were located 1.13 km southeast and 2.5 km northwest of Liubasha.

The kurgan originally consisted of two separate mounds that were subsequently merged into one kurgan. The presumed main burial (№19) in one of the two primary mounds of Liubasha is dated to 2864-2350 calBCE, while burial 16, located under the other primary mound is dated to 3074-2888 calBCE, suggesting that the latter mound containing burial 16 was erected prior to the mound containing burial 19.

Based on the available $^{14}C$ dates, the Liubasha kurgan was in use for at least 1111 ±157 years (3074-2888 calBCE, L16, to 2120-1620 BCE for L9/L12). An undated Scytho-Sarmathian burial excavated in the kurgan[2] extends the use of the Liubasha kurgan potentially another 1500 years.

The kurgan contained a total of 19 burials. Burials 4 (Catacomb archaeological complex), 7, 8, 16 (Yamna archaeological complex), as well as 9 and 12 (identical twins, the Babyne archaeological group) are presented in this report.

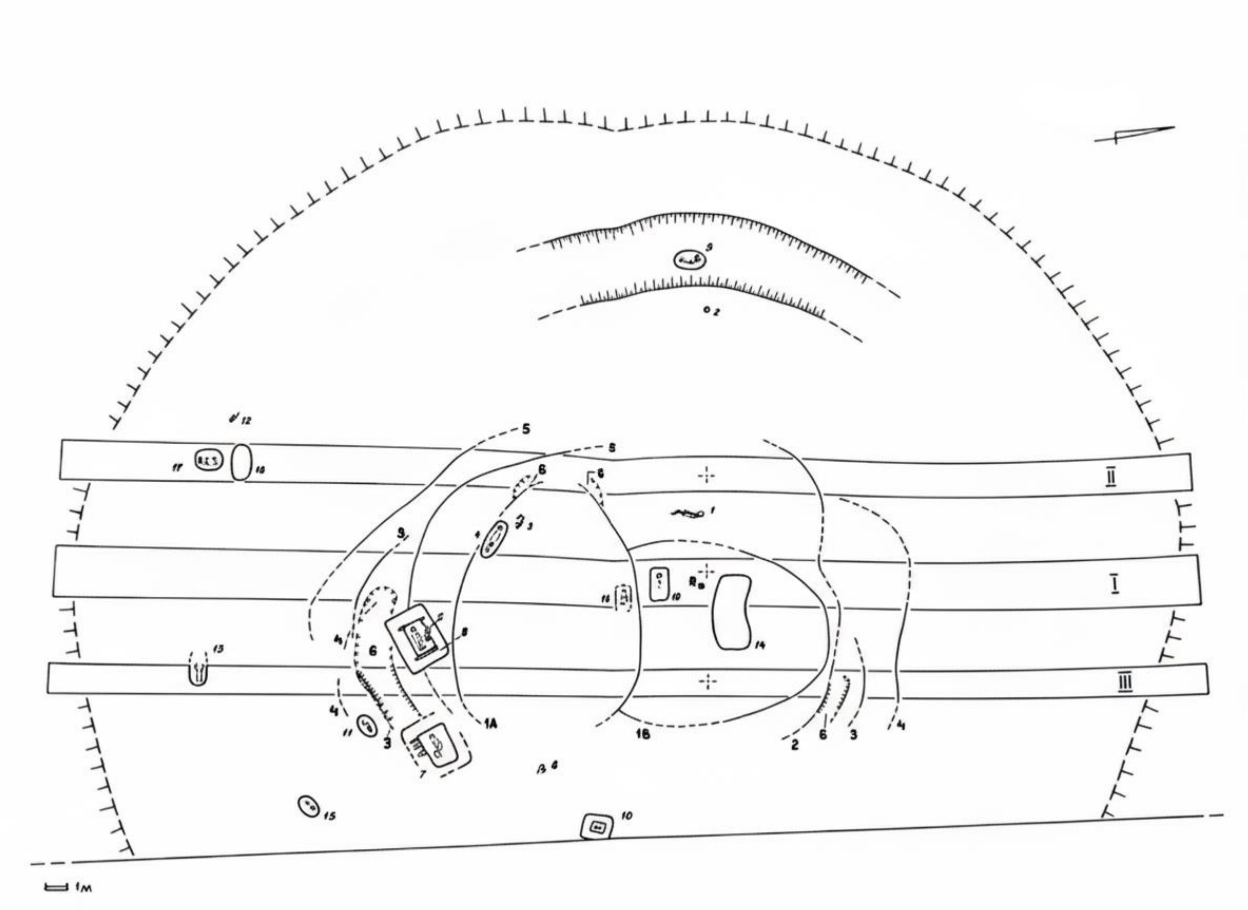

*Figure S6. Kurgan Liubasha excavation plan. Image by S. Ivanova.*

#### Liubasha Kurgan burial 4 (L4)

#### Adult, 2600-2100 BCE

Burial 4 was found at a depth of 2.2 m, 11.2 m to the south-southwest of the benchmark. At this level, a 0.35 × 1.75 m tree trunk was lying in the direction of the north-west - south-east on the edge of the burial. In the central part of the log there was a hollow or hollowed-out hole 0.73 m long, at the bottom of which there was dense greenish clay. The trunk was placed along the southern wall of the burial chamber, on its side, overhanging the pit. The eastern end of the chamber had a width of 0.63 m. The length of the burial chamber was 1.97 m; the depth was 0.4 m (total depth: 2.55 m). The interred lay extended supine along the axis of the pit, with a slight inclination to the right, facing south. The hands were stretched along the body; the left hand was tucked under the pelvis. The skeleton was partially destroyed. A bone fragment was found in the right hand; a fragment of the same bone was found in the tree trunk hollow. White organic decay was traced around the skeleton and at the bottom under the bones, resting on a 2-3 cm layer of chalk.

##### Liubasha Kurgan burial 7 (L7)

##### Adult, 3300-2500 BCE

Burial 7 was found 15.2 m to the southeast of the benchmark, at a depth of 3 m, on the remains of a transverse wooden floor on the ledges of the burial chamber. The upper part of the pit had dimensions of 3 × 2.8 m. The ceiling, consisting of planks 20 cm wide and 2–3 cm thick, sank inside the burial chamber together with mixed filling (loam and chernozem). Near its northern corner, a layer of clay coating 0.45 × 0.32 m, reinforced with twigs, lay on the slabs with the smooth side down. In plan, the burial chamber had the appearance of an oblique trapezium with rounded corners, with a length from the top of 1.8 to 2.0 m; 1.05–1.35 m wide. Towards the bottom, the pit narrowed, reaching dimensions of 1.6 × 1.02 m. The depth of the chamber from the level of the ledge is 0.9 m (total depth: 3.9 m). The interred was laid contracted on back, with the head to the southwest. The arms were extended along the body, while the left hand was brought under the pelvis. The legs, originally placed with the knees up, fell to the right. Dark cherry-colored ocher was noted on the parietal bones, the left and right temporal bones, and the bones of the feet. Two-layered brown-white organic decay was found at the bottom of the chamber. Imprints of plant stems and structural fibers were observed along the left arm in the white rot. Along the right-hand side was a layer of chalk covered by a deposit of dark ocher. Light brown rust was discovered under the pelvic bones.

##### Liubasha Kurgan burial 8 (L8)

##### Adult, 2855-2291 calBCE (3990±70 BP, Ki-11177) [2]

Burial 8 was discovered 14.37 m south-southeast of the benchmark at a depth of 3.25 m. The upper part of a rectangular pit (2.73 × 2.2 m) with slightly curved walls was traced for 0.55 m. A rectangular burial chamber (1.55 × 0.80 m), 1 m deep from the ledge (4.75 m), had been dug into the base of the pit and covered with a wooden flooring composed of 14 transverse planks 0.18–0.20 m wide and averaging 1.55 m in length (range 1.30–2.06 m). The walls of the chamber were likely lined with reed mats, evidenced by a 5–15 mm layer of white dust formed from tightly packed stems. The deceased lay contracted on the back with the head oriented southwest. The head and shoulders were originally elevated, causing the skull to appear drawn into the shoulders and producing a characteristic S-shaped curve in the spine, suggesting the body rested on an organic "pillow." The arms appear to have supported the upper body: the right arm was slightly bent, the left extended. The flexed legs had twisted or fallen to the right side. The left hand rested in a scatter of ocher. An oval patch of crimson ocher was present on the frontal bone, accompanied by small deposits of white organic material. Ocher also outlined the margins of the eye orbits, with a concentration in the left orbit and additional spots on the left side of the cranial vault. White organic material was found beneath and around the skeleton, particularly beneath the head (possibly the remains of the "pillow"), with a layer of brown matter noted beneath it.

##### Liubasha Kurgan burial 9 (L9)

##### Male, adult, 2100-1600 BCE

Burial 9 was located 16.3 m west of the kurgan's reference benchmark at a depth of 4.3 m, within a ditch traced along the western side of the mound and sealed beneath dense silty deposits. The burial pit was oval in plan (1.56 × 0.93 m), trough-shaped in cross-section, and shallow, with a depth of 0.15 m (base at −4.45 m), oriented along a north–south axis. The fill consisted of compact silty sediment. The skeleton was positioned next to the eastern wall of the pit. The individual was interred in a flexed position on the left side, with the head oriented southward. The arms were extended along the body, with the hands joined and resting against the left thigh.

##### Liubasha Kurgan burial 12 (L12)

##### Male, adolescent, ca. 14 yo., 2119-1624 calBCE (3520±80 BP, Ki- 11173) [2]

Burial 12 was found 23.8 m south of the central benchmark at a depth of 4.05 m, outside the main circumference of the kurgan and aligned with the outer ditch identified on the western side of the mound. A distinct burial pit was not observed. The skeleton was recovered from clay loam beneath a dense silty layer that filled a ground depression, possibly remnants of the outer ditch traced on the western side of the kurgan. The individual was interred in a tightly crouched position on the left side, with the head oriented toward the southeast. The arms were flexed at the elbows, with the hands positioned close to the face. Genetic analysis indicates that the individuals interred in burials 9 and 12 were identical twins [1].

Liubasha Kurgan burial 16 (L16)

Male, adult, 3074-2888 calBCE (43330±35 BP, PSUAMS-7790). Burial description is provided in [1,2].

#### Mayaky Kurgans 8 and 10 (46.397, 30.272)

Belyaiivskyi District

A detailed description of the Mayaky necropolis is provided in [1,3,4].

Mayaky Kurgan 8 burial 7 (M8.7)

Immature (3-4), 3800-3300 BCE

The grave was rhomboid-shaped (110 × 80 cm) with a depth of 50-80 cm. The remains of a three-to-four-year-old child were found in dense dark brown loamy soil with "white-eye" inclusions, 10 cm above the bottom of the grave. The skeleton lay flexed on its back, with the skull oriented east-northeast (65°). The arms were bent, and the forearms were placed on the rib cage. The legs were turned with knees to the left. The skull faced left with the crown pointing upwards. Stains of red ocher were present on the skull vault and the metaphysis of the left femur. Grave goods consisted of a black-clay figurine with incised ornamentation, a vessel with two tubular lugs on the rim, and a pot with cord impressions and flat stamp marks containing an Unio shell valve with red ocher residue inside. The vessels were discovered at the level of the skeleton, while the figurine was found at the bottom of the grave.

Mayaky Kurgan 10 burial 2 skeleton 2 (M10.2.2)

Male (50-55), 4336-4065 calBCE (not adjusted for reservoir effect) (5390±30 BP, PSUAMS-7845) [1]. Burial description is provided in [1,4].

#### Revova Kurgan Group (47.268, 30.321)

Shyriaivskyi District

In the vicinity of the village of Revova, two kurgans, numbered 3 and 4, were investigated. These kurgans were situated on the eastern bank of the Velykyi Kuialnyk River, at an elevation of 84 m above the river's thalweg. They occupied the upper portion of a southwestern slope that descended towards the river, approximately 1.5

km away from the kurgans. The topographic conditions provided an unobstructed view of kurgan 3 from the south and southwest. Kurgan 3, standing at a mere 1.1 meters tall, was strategically positioned at the convergence of orographic lines. This placement granted it a commanding line of sight extending over 25 kilometers down the Velykyi Kuialnyk River [2,5].

Kurgan 3, 1.1 × 45 m, contained 21 burials, four of them are presented in this report. These include burials 13 (Catacomb), 14 (Babyne), 15 (Yamna), and 19 (Eneolithic, Usatove archaeological group). Based on available $^{14}C$ dates, the Revova 3 Kurgan was in use for 1730±160 years.

##### Revova Kurgan 3 burial 13 (R3.13)
##### Male, 2580-2207 calBCE (3940±60 BP, Ki-11172) [1,2].

Burial 13 was located 8.7 m south-southwest of the benchmark at a depth of 1.9 m. On the natural subsoil, it was visible as the dark fill of a ground-level catacomb with a collapsed vault. The burial chamber had an oval plan measuring 2.1 × 1.5 m and a traced depth of 1.1 m (3.0 m absolute). A rounded entrance shaft adjoined the chamber on the southeast side; it measured 0.7 m in diameter and 0.5 m in traced depth. The deceased lay extended on the back with the head oriented north-northwest. The skull was tilted toward the right shoulder, with the face turned south, and the spine was slightly curved. A ceramic round-bottomed bowl with a neatly cut rim (height 16.5 cm; diameter 18.5 cm) decorated with a band of six grooves had been placed near the eastern wall at the level of the forearms on a bed of burgundy ocher, with its side projections facing south. Three flints were found behind the parietal portion of the skull, and another lay near the left forearm. Each flint was retouched on the ventral surface. Dark brown organic decay was observed on the chamber floor beneath the bedding.

##### Revova Kurgan 3 burial 14 (R3.14)
##### 2139-1748 calBCE (3590±70 BP, Ki-11175) [1,2].

Burial 14 was identified from fragments of a transverse wooden covering, preserved as brown organic decay in the central part of the pit directly above the skeletal remains. The burial was located 6.5 m south of the benchmark at a depth of 1 m. The floor of the burial chamber was traced only partially, based on the brown decay of an organic bedding. The chamber was apparently rectangular with strongly rounded corners, measuring 1.35 m in length and 0.8 m in width. The walls were traced to a height of 0.15 m (1.15 m absolute). The contracted skeleton lay on the left side, oriented west–southwest. The left humerus was parallel to the main axis of the skeleton, while the right humerus was directed toward the left. The remaining arm bones were not preserved. The burial had been sprinkled with bright red ocher, with deposits visible on both the chamber floor and the bones.

##### Revova Kurgan 3 burial 15 (R3.15)

##### Male, 2458-2031 calBCE (3780±70 BP, Ki-11060) [2]

Burial 15 was located 1.5 m south of the central benchmark at a depth of 0.7 m and was sealed beneath a timber covering. The burial pit was rectangular in plan, measuring 1.9 × 1.1 m, with a traced depth of 0.5 m (total depth 1.2 m). The timber covering consisted of massive logs approximately 20 cm in diameter, preserved as fragments along the edges of the pit and within the fill. Along the long sides of the pit floor, narrow grooves approximately 12 cm wide and 2 cm deep were observed. Beneath two of the walls, circular postholes measuring approximately 7 cm in diameter and 19–20 cm in depth were cut into the pit floor. One of these, located near the short eastern wall, contained preserved wood remains. Impressions of closely packed plant stems, 2–5 mm in diameter, were locally preserved on the pit walls. The pit floor was coated with a layer of light-colored clay and sprinkled with ochre. The deceased was placed supine, with the head oriented toward the southwest. The arms were extended along the body, while the legs were flexed and raised at the knees. The vertebral column exhibited curvature in the thoracic region. Above the prepared floor surface, in the area corresponding to the torso, a lens of gray clay was identified, overlain by a uniform white organic residue.

##### Revova Kurgan 3 burial 19 (R3.19)

Male, adult, 3711-3639 calBCE (4905±20 BP, PSUAMS-4763) [1]

Detailed description of the Revova burial 19 can be found in [1,2,5].

**S1 Table. Stable isotopes of carbon and nitrogen used to construct the graph in Fig 3.**

| Population/specimen ID | $^{14}C$ date | $\delta^{13}C$ (‰) | $\delta^{15}N$ (‰) | $\delta^{13}C$ SD | $\delta^{15}N$ SD | Sample size | Ref. |
|---|---|---|---|---|---|---|---|
| West Manych/North Caucasus Eneolithic-EBA | 4300-2700 BCE | -17.40 | 14.80 | 1.07 | 1.57 | 30 | [6–8] |
| Eastern North Pontic Eneolithic-EBA | 4200-2200 BCE | -18.55 | 11.91 | 0.71 | 0.94 | 31 | [9] |
| Western North Pontic Eneolithic-EBA | 3700-2200 BCE | -19.19 | 12.01 | 0.24 | 0.82 | 23 | [1] |
| D1.9 | 2575-2348 calBCE | -19.40 | 12.01 | | | 1 | [1] |
| K1.10 | 3951-3715 calBCE | -18.05 | 15.17 | | | | [1] |
| K1.13 | 3018-2911 calBCE | -19.1 | 12.26 | 0.023 | 0.071 | 1 (2 readings) | [1] |
| K1.21E | 2896-2701 calBCE | -19.5 | 11.35 | | | 1 | This report |
| K2.1 | 3341-3093 calBCE | -19.1 | 12.63 | | | 1 | [1] |
| L9 | 2100-1600 BCE | -17.43 | 14.08 | | | 1 | This report |
| L12 | 2119-1624 calBCE | -18.8 | 13.05 | | | 1 | [2], this report |
| L16 | 3074-2888 calBCE | -19.6 | 11.75 | | | 1 | [1] |
| R3.19 | 3711-3639 calBCE | -19.7 | 11.52 | | | 1 | [1] |
| Taraclia II.2.14 | 3340-3035 calBCE | -17.8 | 13.69 | | | 1 | [1] |
